# Coordinated downregulation of the photosynthetic apparatus as a protective mechanism against UV exposure in the diatom *Corethron hystrix*

**DOI:** 10.1101/252668

**Authors:** Robert W. Read, David C. Vuono, Iva Neveux, Carl Staub, Joseph J. Grzymski

## Abstract

The effect of ultraviolet radiation (UVR) on photosynthetic efficiency and the resulting mechanisms against UV exposure employed by phytoplankton are not completely understood. To address this knowledge gap, we developed a novel close-coupled, wavelength-configurable platform designed to produce precise and repeatable *in vitro* irradiation of *Corethron hystrix*, a member of a genera found abundantly in the Southern Ocean where UV exposure is high. We aimed to determine its metabolic, protective, mutative, and repair mechanisms as a function of varying levels of specific electromagnetic energy. Our results show that the physiological responses to each energy level of UV have a negative linear decrease in the photosynthetic efficiency of photosystem II proportional to UV intensity, corresponding to a large increase in the turnover time of quinone re-oxidation. Gene expression changes of photosystem II related reaction center proteins D1, CP43 and CP47 showed coordinated downregulation whereas the central metabolic pathway demonstrated mixed expression of up and downregulated transcripts after UVR exposure. These results suggest that while UVR may damage photosynthetic machinery, oxidative damage may limit production of new photosynthetic and electron transport complexes as a result of UVR exposure.

## Introduction

Diatoms are microscopic photosynthetic algae that are ubiquitous throughout the surface waters of the oceans (Lohman 1960), account for roughly 40% of oceanic primary production, and one-fifth of the Earth’s total primary production (Falkowski and Raven 2007). Diatoms also play a vital role in the global carbon cycle through their uptake of dissolved CO_2_ and subsequent carbon fixation that forms the base of the marine food web (Armbrust 2009). Given their global distribution, diatoms have adapted to survive under a variety of environmental conditions (Ligowski et al. 2012; Verde and Prisco 2012; Marchetti et al. 2012). Unfavorable conditions such as nutrient limitation (Allen et al. 2008; Dyhrman et al. 2012; Shrestha et al. 2012; Bender et al. 2014), varying light levels (Domingues et al. 2012; Herbstová et al. 2015) and UV exposure (Wu et al. 2015) are commonplace. The latter is of particular interest because of the damaging effects UV can have on photosynthesis as well as other metabolic pathways.

In phytoplankton, UV exposure can inhibit photosynthesis, based on the relative dose and dose rate (Cullen and Lesser 1991). An inhibition of the photosynthetic rate causes a decrease in the rate of primary production, with consequences in marine ecosystems as well as terrestrial environments. Phytoplankton can also produce protective compounds to combat the deleterious effects of UVR such as mycosporine-like amino acids (MAAs), DNA photolyases and many more undefined compounds (Helbling et al. 1996; Coesel et al. 2009). These compounds, when added to commercial products such as sunscreen (Berardesca et al. 2012; Emanuele et al. 2013), show promise in reducing carcinogenic effects of UV exposure in humans. Advances in this field require a reliable method to induce and measure damage in a controlled laboratory setting.

Fluorescence kinetics measurements are a reliable estimator of photosynthetic electron transport rates and photosystem II health (PSII) (Kolber and Falkowski 1993) in photoautotrophic organisms. Because PSII is a target of UVR induced damage (Tevini and Teramura 1989; Szilárd et al. 2007), Fast Repetition Rate Fluorometry (FRRF) has been used to examine variations in several photosynthetic parameters in relation to light impacts on the cell (Kolber et al. 1998). Changes in parameters such as the maximum quantum yield of PSII (Fv/Fm) (Geider et al. 1993; Kolber and Falkowski 1993), the turnover time of electron transport from QA -> QB (tau1) and QB -> PQ (tau2) (Kolber et al. 1988), and the functional cross section of PSII (σ _PSII_) -- the effective target size of the PSII antenna in Å^2^ (quanta)^−1^ (Kolber et al. 1998), act as proxies to monitor electron transport rates and the relative health of PSII. Here we use them to monitor the rate and intensity of photosynthetic damage within the cell. Changes in these parameters are a function of the dose and dosage rate of absorbed radiation, as this has a direct impact of the oxidation state of PSII electron transport chain.

In this study, the diatom *C. hystrix* (CCMP 308) was subjected to increasing intensities of UVR energy ranging from 0.32 mW/cm^2^ to 1.59 mW/cm^2^ using a custom built UVR emitter array. This gave us the ability to precisely control and measure damage as a decrease in the photochemical efficiency of PSII using FRRF. Fv/Fm, sigma/σ _PSII_, and tau were monitored hourly or bihourly to measure the UV damage to PSII relative to non-irradiated conditions. Our aim was to test the UVR emitter array to identify break points in photochemical efficiency, as measured by FRRF, to better understand the physiological constraints of an ecologically important phytoplankton to UVR exposure. Furthermore, we aimed to characterize the transcriptomic profile of *C. hystrix* under 0.64 mW/cm^2^ UVR intensity to reveal transcriptional responses in the central metabolic pathway, photosynthetic electron transport chain (ETC), and DNA repair mechanisms. This study thereby provides a comprehensive investigation of the physiological and molecular stress response to UV irradiation using a UV emitter array designed specifically to dose planktonic phototroph cultures with any desired UVR intensity. These tools and methods, if shown to be successful, could be used to manipulate organisms to better understand how they protect themselves against DNA damage.

## Materials and Methods

### Cell Cultures

*Corethron hystrix* CCMP 308 (Bigelow Laboratory for Ocean Sciences, East Boothbay, Maine, USA) was grown at a maintenance temperature of 14 °C under 12:12 L:D at an illumination of ~40 μmol photons m^−2^ s^−1^ using white LEDs. Although 14 °C was the recommended temperature for *in vitro* studies, *C. hystrix* has a much wider known temperature range. Duplicate cultures were grown in L1 medium (Guillard and Hargraves 1993), prepared with 0.2 μm filtered surface seawater from the Gulf of Maine (Bigelow Laboratory for Ocean Sciences, East Boothbay, Maine, USA). Chlorophyll pigment was extracted in 90% acetone at −20 °C for 17 hours, in the dark. Following extraction, fluorescence of each sub-sample was measured using a 10AU Fluorometer (Turner Designs, Sunnyvale, CA, USA), and chlorophyll concentrations were calculated. Cell counts were determined using a Sedgewick chamber under bright light. Specific growth rates of culture replicates were estimated from the growth curves constructed from chlorophyll *a* fluorescence and cell counts obtained under non-irradiating conditions. Growth rate (units of doublings per day) was calculated from log-normalized exponential growth phase. Growth curves were used to determine the mid-exponential phase when UVR irradiation would be performed. During the UVR experiments, chlorophyll *a* measurements and morphological cell counts were also collected bihourly in order to changes affected by UVR.

### Photosynthetic Kinetics

Fv/Fm, sigma, and tau measurements were monitored using a FRRF (Soliense, Inc, Shoreham, NY, USA). Cell cultures, in biological duplicate, were subjected to either bright white light only at ~ 40 μmol m^−2^ sec^−1^ (control condition) or a combination of bright white light (~ 40 μmol m^−2^ sec^−1^) and UVR exposure ranging from 0.32 mW/cm^2^ to 1.59 mW/cm^2^ (experimental condition). UVR was performed using an LED emitter platform (Lumenautix Inc., Reno, NV, USA; Supplementary Fig. S1). The overall experimental period consisted of both control and experimental conditions running for six hours, followed by a dark period of six hours for the two conditions.

### NEST Configurable Emitter Array

The NEST array is comprised of multiple discrete solid-state emitters operating in two modes, VIS and UV connected to a control unit (Supplemental Text). The output is regulated and temperature compensated for constant current DC – allowing for precise control and stability of light output. Visible LEDs are blue (3), red (3) and white (6) and the UV emitters (3). For this work only the UV and white LEDs were used. The white LED color temperature is 3985K and the output can vary between 4 and 380 μmol m^−2^ sec^−1^.

The LED array allows for the precise and repeatable incremental *in vitro* irradiation of target organisms to determine their protective, mutative, and repair properties as a function of varying levels of specific electromagnetic energy (Supplementary Fig. S1). The energy categories can be adjusted incrementally and independently to affect the organism’s biological functions and to stress the organism to evoke specific physiological responses or cause DNA / RNA mutations.

Net UVR organism exposure was reported from the emitter array’s gross output after subtracting attenuation from surface reflection, transmittance, absorption, and diffusion of the quartz glass container and seawater. This attenuation was approximately 25% of the gross emitter output. Data were collected every hour for UVR energy intensities of 0.32 and 0.64 mW/cm^2^ (measured at 285 nm, 12 nm FWHM). At higher doses of 0.96 – 1.59 mW/cm^2^ (285 nm 12 nm FWHM), measurements were collected every 30 minutes. Raw data were processed and plotted using the ggplot2 package in the R environment (Wickham 2009). Rate constants for both Fv/Fm and sigma were calculated using a linear regression model also using R.

### Illumina sequencing and Gene Expression analysis

After photosynthetic kinetic measurements were made, total RNA was extracted immediately in duplicate from cells in the mid-exponential phase of growth directly after UVR exposure (0.64 mW/cm^2^, experimental and control cultures), as well as directly after a dark recovery period of six hours (experimental and control cultures), using the Ambion ToTALLY RNA kit (Life Technologies, Grand Island, NY, USA). The same experimental and control cultures in duplicate were used during both harvesting periods. These extractions produced approximately ~10-12 μg of total RNA from each pellet. Samples were sent for sequencing at the Biodesign Institute at Arizona State University (Tempe, AZ, USA). Library preparation was fully automated and performed using the Apollo 324 liquid handling platform with selection for polyA RNA. Illumina HiSeq Sequencing yielded 2×100 bp paired-end reads.

Raw sequencing reads were uploaded to the sequencing read archive (NCBI accession: SRP091884, SRX2255404) and inspected using Fastqc (Andrews 2009) to determine quality, ambiguous read percentage and relative amount of sequence reads. Illumina RNA sequencing resulted in an average of 19.4 million raw reads per cDNA library with an average quality score of Q38 (Supplementary Table S1).

Raw Sequencing reads were trimmed using the sequence trimming program Trimmomatic with the following options: Remove any Illumina adapter, cut off the end of any read where the quality score falls below 10, use a sliding window of 5 to cut and trim any base where the average quality score falls below 32 for that window and only keep trimmed reads with a minimum length of 72. (Bolger et al. 2014). De-novo transcript assembly was performed using Velvet (Zerbino and Birney 2008). Optimal k-mer selection, as well as read coverage cutoff selection, was determined by the VelvetOptimiser (Gladman and Seemann 2012). Velvet-constructed contigs were assembled into full-length transcripts using the Oases transcriptome assembler, with a minimum transcript length of 150 base pairs (Schulz et al. 2012). Raw sequences were aligned to the assembled transcripts by Bowtie2 (Langmead and Salzberg 2012). Abundances of mapped sequence reads were calculated using eXpress (Roberts and Pachter 2012), providing the estimated count of reads that mapped to each individual assembled transcript. Estimated counts from eXpress were normalized and counts were calculated using DESeq2 (Love et al. 2014). Transcripts were considered differentially expressed if their associated log_2_ fold changes were significant at (adjusted) *p*<0.05, based on the Wald test of DESeq2, while controlling for false discovery using the Benjamini-Hochberg Procedure (Love et al. 2014). Based on our experimental design, differential expression was compared between the UVR+white light irradiated cells (experimental treatment – in duplicate) and bright white light only cells (control treatment – in duplicate), directly after the six hour UVR exposure ended and also after a six-hour dark recovery period.

*C. hystrix* is not well annotated. In order to artificially reconstruct the pathways in *C. hystrix*, transcripts from our de-novo assembly were *in silico* translated and compared to Uniprot proteins using Hhblits, which is part of the HH-suite software package (Remmert et al. 2011; The UniProt Consortium 2017). Homologous protein homology was inferred from *in silico* translations using hidden Markov model alignments from Hhblits (Soding 2005). Translated transcripts were considered homologs if there was a >90% probability of the translated transcript being a homolog to a Uniprot protein.

Subsequent classification of homologous Uniprot proteins into functional annotation groups was performed by grouping Uniprot Ids based on gene ontology (Ashburner et al. 2000; The Gene Ontology Consortium 2017). In the case of supplemental photosynthetic electron transport, gene ontology provided a poor representation of the selected pathway. For this pathway, homologous proteins from the model centric diatom *Thalassiosira pseudonana* were combined with the Uniprot annotations, with the understanding that this organism’s annotations may have changed since they were first annotated and uploaded to public databases in 2004 (Armbrust et al. 2004). The entire annotated *Thalassiosira pseudonana* photosynthetic electron transport pathway was downloaded from the Kyoto Encyclopedia of Genes and Genomes (KEGG) (Kanehisa and Goto 2000; Kanehisa et al. 2013).

Translated transcripts mapping to a homolog were sometimes one of several contigs. To maintain consistent mapping between homologs, transcript contigs were binned by functional annotation after translation. Additionally, in most cases, up and downregulation variation between contigs was small, making expression patterns more evident. However, on occasion, transcripts for the same functional annotation but different loci were both up and downregulated. Because *C. hystrix* does not have an annotated genome, binning by functional annotation produces an expression overview for the homolog, which accounts for transcriptional variability. Binned isoforms were visualized with a box and whisker plot using the ggplot2 package (Wickham 2009). The line in the box represents the median log_2_ fold change when combining all the isoforms. The hinges are the 1^st^ and 3^rd^ quartile.

## Results

### Chlorophyll A and Cell Counts under laboratory conditions

*C. hystrix* grew at rate of 0.37 doublings per day based on chlorophyll *a* concentration (linear regression equation y=5.74 + 0.254x, r^2^=0.991) (Supplementary Fig. S2a); this was consistent with microscopic cell counts, which produced a growth rate of 0.392 doublings per day (linear regression equation y=9.70+0.272x, r^2^ = 0.990) (Supplementary Fig. S2b). Under UV light of different intensities, chlorophyll *a* content remained relatively constant during the six-hour irradiation period (Supplementary Fig. S3). Additionally, there was no significant increase in cell growth over the irradiation period. However, light microscopy revealed morphologically altered cells in the UVR exposed cultures compared to control cultures. The percentage of morphologically intact cells was mostly constant for the lower UVR intensity levels (0.32 – 0.64 mW/cm^2^), with a small decrease during the last two hours at 0.64 mW/cm^2^ (Fig. 1). However, higher intensities (0.96 – 1.59 mW/cm^2^) caused a pronounced decrease in the amount of morphologically intact cells, especially during the last two hours of UVR exposure (Fig. 1).

**Figure 1:**
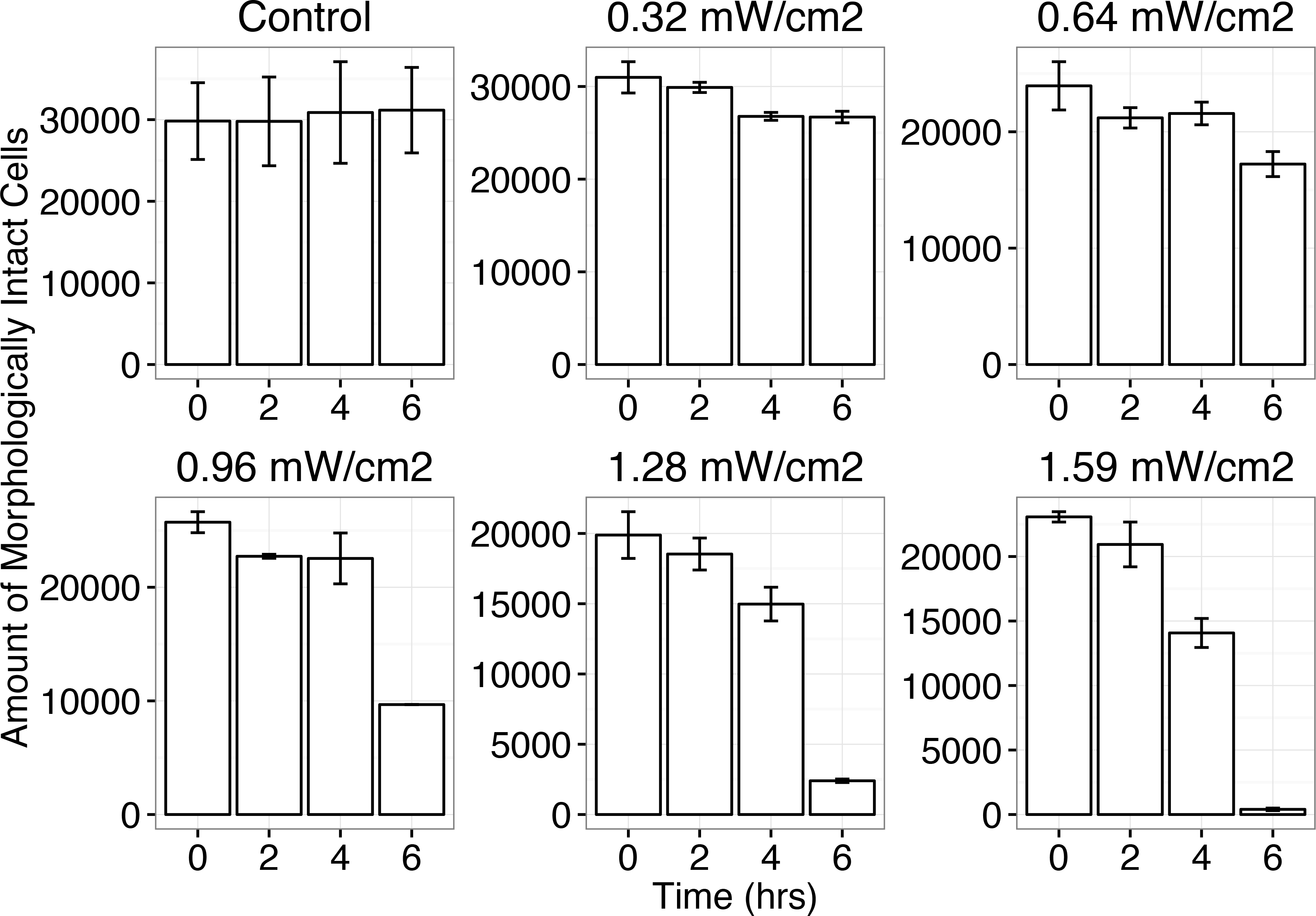
Morphologically intact cells during the 6 hour UVB irradiation period. Morphologically intact cells were estimated by bright light microscopy over the 6-hour irradiation period. All counts were collected both the non-irradiated cells and cells under 5 different intensities of UVB (0.32 mW/cm^2^ – 1.59 mW/cm^2^). Intact cell counts were collected in triplicate every 2 hours with error bars representing the standard deviation between replicated counts.

### Photosystem II

To monitor the changes in PSII during exposure to UVR, photosynthetic kinetic measurements were recorded for *C. hystrix* using FRRF (Fig. 2). Several photosynthetic parameters were derived from FRRF measurements. First, Fv/Fm empirically represents the maximum quantum yield of PSII (Fig. 2A) (Geider et al. 1993; Kolber and Falkowski 1993). Fv/Fm is a dimensionless parameter representing how efficiently absorbed photons are used for electron flow: a value of 1 represents complete absorbance efficiency and 0 represents no absorbance (Suggett et al. 2009). A second parameter, sigma is a proxy for the product of the optical cross section for PSII (roughly proportional to the number of chlorophyll molecules per PSII), the efficiency of excitation transfer from the antenna to the PSII reaction center and the quantum yield of charge separation (Fig. 2B) (Mauzerall 1986; Oxborough et al. 2012).

**Figure 2:**
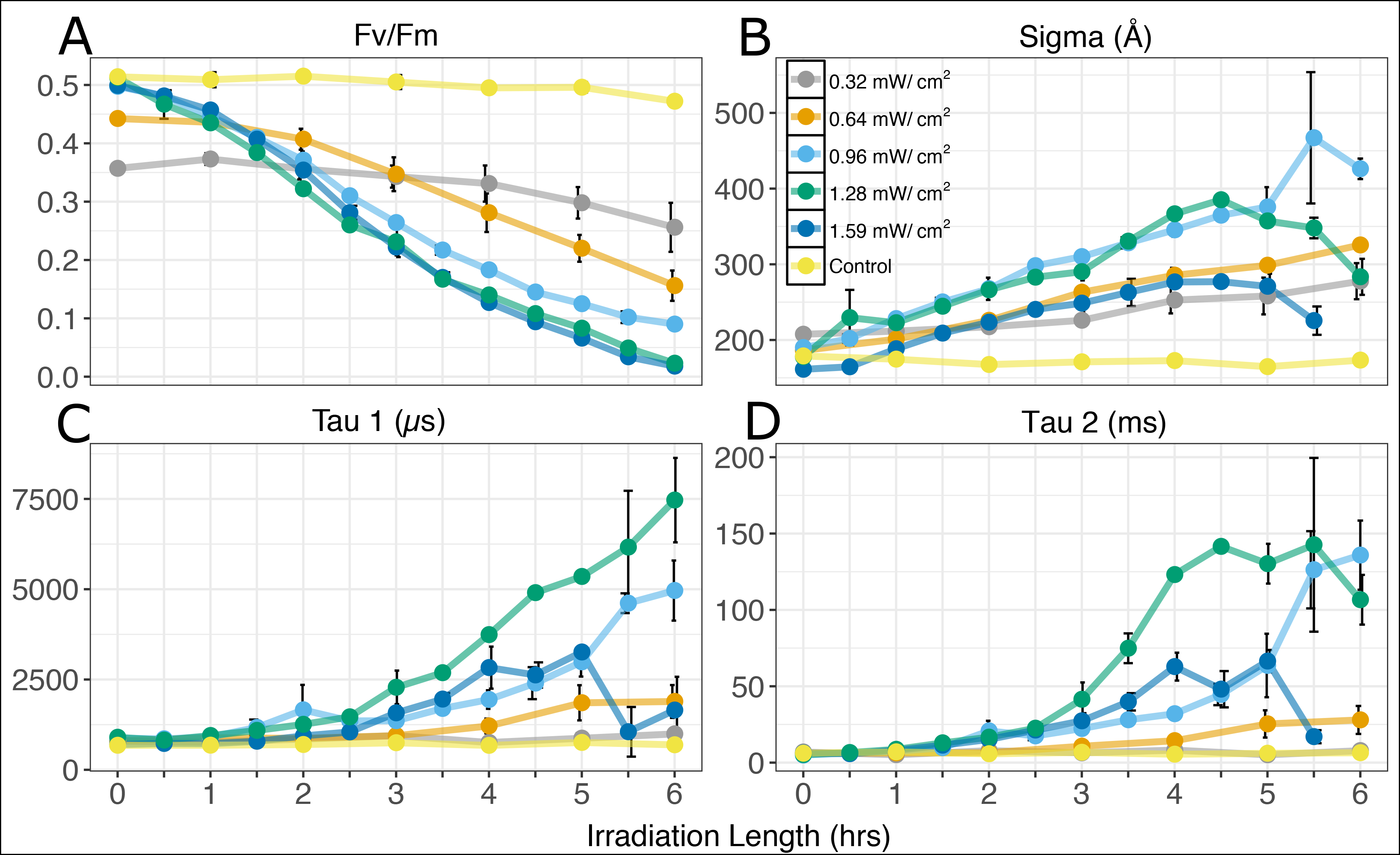
**A:** Fv/Fm as a measure of the maximum quantum yield of PSII. **B:** Sigma as a proxy for functional cross section and effective target size of the PSII antenna in Å^2^ (quanta)^−1^. **C**: Tau 1 turnover time. During every hour at 0.32 mW/cm^2^ and 0.64 mW/cm^2^, and every half hour at 0.96-1.59 mW/cm^2^ the tau 1 turnover time was calculated using Fast Repetition Rate Fluorometry (FRRF). Irradiation time is on the x-axis with the turnover time in μs on the y-axis. **D**: Tau 2 turnover time. During every hour at 0.32 mW/cm^2^ and 0.64 mW/cm^2^, and every half hour at 0.96-1.59 mW/cm^2^ the tau 2 turnover time was calculated using Fast Repetition Rate Fluorometry (FRRF). Irradiation time is on the x-axis with the turnover time in μs on the y-axis. Note the drop in turnover time at 1.28 and 1.59 mW/cm^2^.

Fv/Fm decreased linearly (Supplementary Table S2) with the rate of decay increasing with intensity (Supplementary Fig. S4-S8, Supplementary Tables S3-S7). Conversely, sigma increased linearly with increasing intensity, demonstrating an inverse relationship between sigma and Fv/Fm for all doses of UVR in this study, with sigma increasing and Fv/Fm decreasing as damage accumulated (Fig. 2 AB, Supplementary Fig. S4-S8). At the higher UVR intensities there was a notable decrease in sigma as photosynthesis became inhibited (Supplementary Table S2). Overall, UVR had a powerful effect on the fluorescence kinetics of PSII, especially at the higher UVR intensities (0.96 – 1.59 mW/cm^2^) (Fig. 2, Supplementary Tables S5-S7). For example, sigma increased only 15% from the start of the UVR exposure until the end for the lowest UVR intensity of 0.32 mW/cm^2^, with the largest increase evident after three hours under irradiation (Supplementary Fig. S4, Supplementary Table S3). In comparison, at 0.96 mW/cm^2^, we observed a similar 15% increase in sigma within the first two hours of the start UVR exposure, with a total increase in sigma of approximately 56% over the irradiation period (Supplementary Fig. S6, Supplementary Table S5). Furthermore, UVR intensities of 0.96 mW/cm^2^ and 1.28 mW/cm^2^ had very strong and similar responses. The rate of change of sigma for each of these two treatments was almost identical and approximately 44% faster than the rate of change observed at 0.64 mW/cm^2^ (Supplementary Table S2) and 72% faster than the rate observed at 0.32 mW/cm^2^. At the highest intensity of 1.59 mW/cm^2^, the rate of change for sigma was slower than the rate at 0.96 mW/cm^2^ and 1.28 mW/cm^2^, likely because of extreme damage to the photosynthetic reaction centers.

We used RNA-seq to evaluate the expression response of *C. hystrix* PSII transcripts for a single UVR intensity (0.64 mW/cm^2^). This intensity was chosen because the majority of cells remained morphologically intact throughout the six-hour irradiation period (Fig. 1). There were 12 differentially expressed PSII related transcripts detected directly after UVR exposure (Fig. 3, Supplementary Fig. S9). All 12 transcripts were downregulated and demonstrated little variation in their fold changes, indicating a strong and coordinated transcriptional response (Fig. 3, Supplementary Fig. S9, Supplementary Table S8). Specific transcripts, related to proteins such as the reaction center core D1 and D2 proteins were significantly downregulated (adjusted *p* < 0.001; Supplementary Table S8), corresponding with observed decreases in the photosynthetic efficiency. Moreover, translated transcripts mapping to cytochrome c550, cytochrome b559 and PSII PsbH proteins were also strongly downregulated.

**Figure 3:**
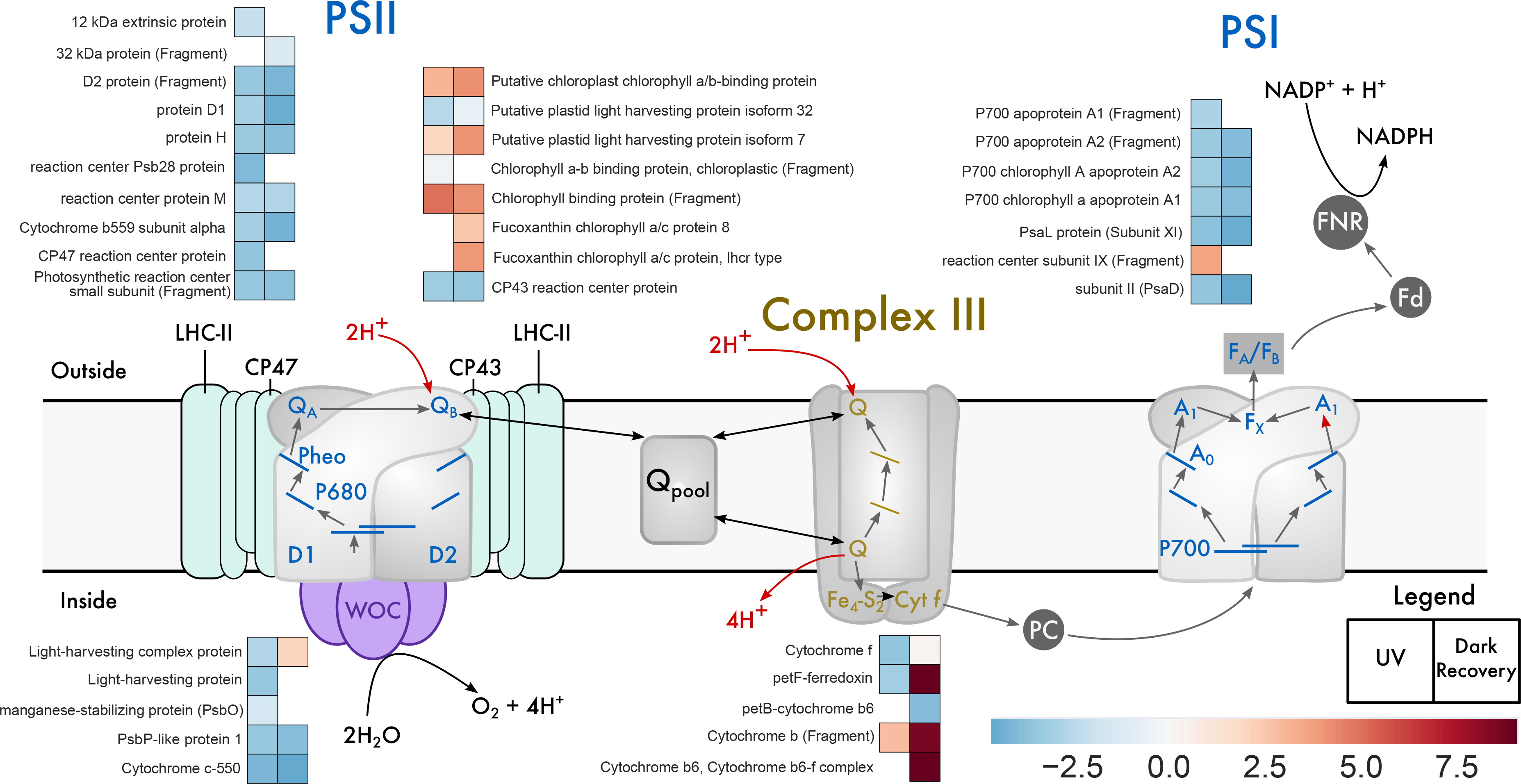
Metabolic pathway reconstruction of the photosynthetic electron transport chain. Uniprot names for each gene product correspond to heat map subplots in the order shown (left-to-right for condition (directly after UVR and after dark recovery) and top-to-bottom for each Uniprot name). Higher transcript abundance is represented in red (upregulation), lower transcript abundance in blue (downregulation) and empty spots represent a Uniprot homolog wasn’t differentially expressed during that condition.

### Reoxidation of the Electron Acceptors

There was very little change in tau 1 during the first 2.5 hours for any dose of UVR (Fig. 2C). After 2.5 hours, turnover time began to increase for the larger UVR doses: 0.96, 1.28, and 1.59 mW/cm^2^ (Fig. 2). Overall, tau 1 turnover time increased from 26.61 μs at 0.32 mW/cm^2^ to 1114.0 μs at 1.28 mW/cm^2^ – an approximately 42x increase (Supplementary Table S9). At the highest UVR energy (1.59 mW/cm^2^), tau 1 turnover time initially followed the same trend as 1.28 mW/cm^2^. However, there was a significant decline in the turnover time after 5 hours (Fig. 2C, Supplementary Table S9), likely because of a severe inhibition in photosynthesis.

Tau 2 reacted similarly to tau 1 during UVR exposure, although turnover time for tau 2 was much longer, on the order of milliseconds instead of microseconds (Fig. 2D). For the first two hours of irradiation, there was no significant change in the turnover rate for tau 2 (Fig. 2D). After 2.5 hours, tau 2 increased with each successive increase in UVR energy, with the exception of 1.59 mW/cm^2^, which decreased after 5 hours, similar to the behavior of tau 1 for the same treatment (Fig. 2D, Supplementary Table S10).

Photosynthetic electron transport genes are not well annotated in the Uniprot database. Thus, proteins from the ubiquitous model centric diatom *T. pseudonana* were combined with available Uniprot proteins to test for electron transport protein homology, as the majority of the photosynthetic electron transport pathway is partially annotated by KEGG (Kanehisa and Goto 2000; Kanehisa et al. 2013). Comparative transcriptomics for the supplemental electron transport pathway of the Z scheme at 0.64 mW/cm^2^ were similar to the observations of PSII gene expression. There were 10 transcripts related to supplemental electron transport, most of which are a subunit of the cytochrome b_6_f complex. Cytochrome f (*petA*), the largest subunit of the cytochrome b_6_f complex (Gray 1992), was downregulated according to both our *T. pseudonana* custom database and Uniprot database. Differential expression analysis of the 10 transcripts show nine decreased in abundance and one increased, demonstrating a coordinated downregulation of the supplemental electron transport genes. These results are corroborated by the increases in tau 1 and tau 2 turnover times (0.64 mW/cm^2^).

### Photosystem I and RuBisCO

Photosynthetic kinetic measurements for photosystem I (PSI) are not reported in this study because they are outside the resolution of the current FRRF machine. However, gene expression of PSI genes was analyzed for the 0.64 mW/cm^2^ UVR treatment (Fig. 3). There were 16 PSI related transcripts; 15 transcripts decreased in abundance compared to non-irradiated samples after UVR exposure. Both *psaA* (Apoprotein A1) and *psaB* (Apoprotein A2), which bind through hydrophobic interactions to form the core complex of PSI (Falkowski and Raven 2007) were downregulated under UVR. Additionally, psaD (PSI subunit II) transcripts, which facilitate the docking of ferredoxin and are an essential component of correct PSI function, were downregulated (Hou et al. 2017). Inorganic carbon fixation involves the ribulose-1,5-bisphosphate carboxylase/oxygenase enzyme (RuBisCO). Five transcripts that map to RuBisCO related proteins were downregulated compared to the non-irradiated samples.

### Light Harvesting Complex

Light harvesting complexes are photosynthetic protein complexes that harvest light energy and channel it into the PSII and PSI reaction centers. In contrast to the downregulation of PSII and PSI proteins, several light harvesting complex transcripts were upregulated in response to UVR. There was a total of 39 transcripts that mapped to light harvesting related proteins (Fig. 3, Supplementary Table S8). Approximately 54% (21/39) of those transcripts were decreased in abundance compared to the non-irradiated samples, including the highly conserved PSII CP43 and CP47 proteins.

### Metabolic Pathway Expression

Glycolytic regulation in plants is accomplished by three main proteins: hexokinase, phosphofructokinase and pyruvate kinase (Plaxton 1996). From the glycolysis pathway, there were 24 differentially expressed transcripts mapping to eight homologous Uniprot proteins. In general, glycolytic Uniprot homologous transcripts showed mixed expression. Three *in silico* translated transcripts mapping to homologous proteins phosphofructokinase, phosphoglycerate kinase and enolase decreased in abundance directly after UVR exposure. Five *in silico* transcripts mapping to homologous proteins aldolase, triosephosphate isomerase, G3P dehydrogenase, phosphoglycerate mutase, and pyruvate kinase increased in abundance directly after UVR exposure (Fig. 4, Supplementary Table S11). None of the transcripts in our data mapped to hexokinase homologs based on the results from both HMM protein detection and BLASTX (Camacho et al. 2009).

**Figure 4:**
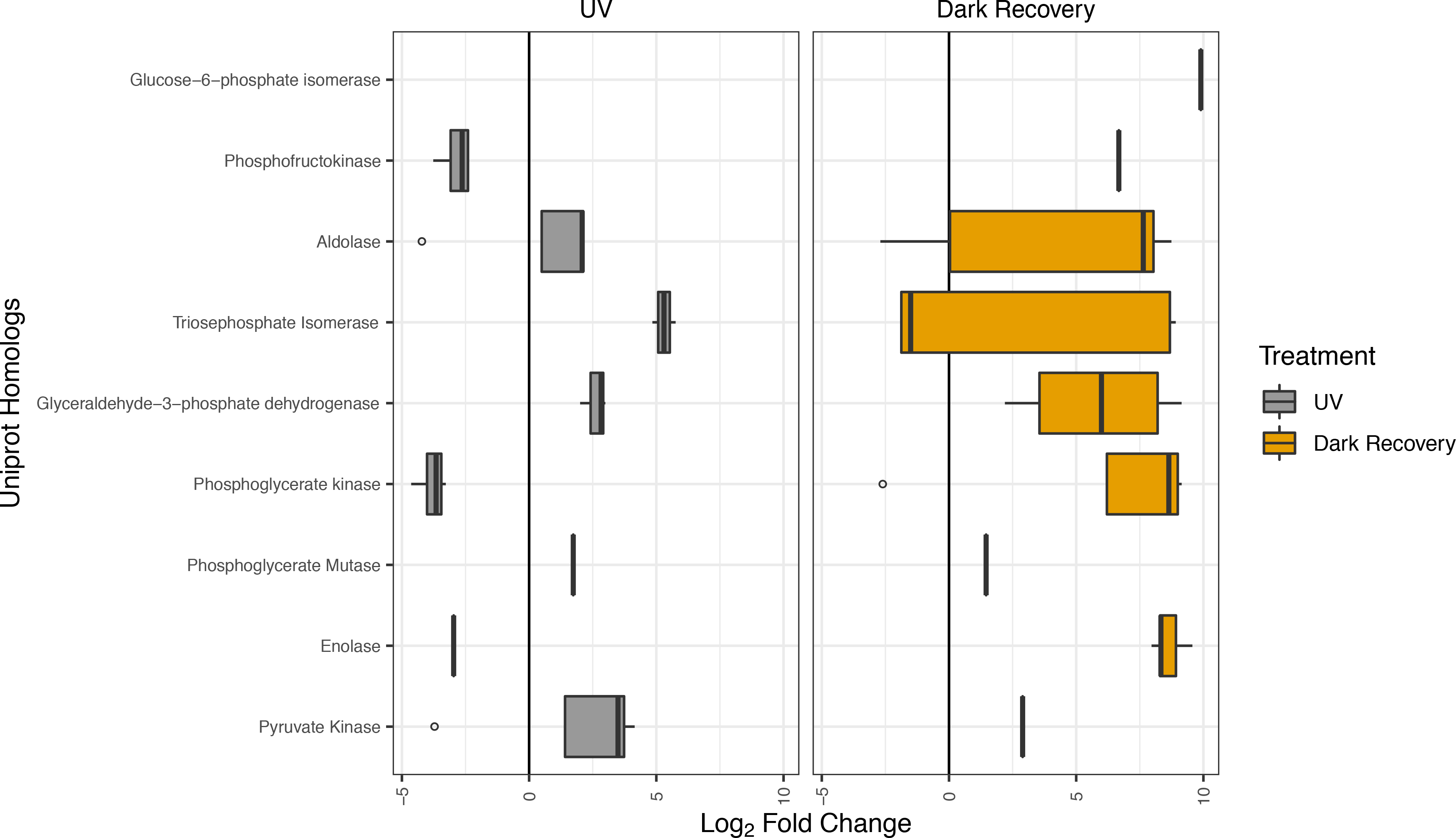
Box and whisker plot for Uniprot glycolysis homologs. Log_2_ fold changes directly after UVR irradiation are represented in the left figure – labeled “Day” – while log_2_ fold changes after six-hour dark recovery – labeled “Night” – are represented in the figure to the right. The line within the box is the median of the log_2_ fold changes for that specific homolog. The hinges are the 1^st^ and 3^rd^ quartile. The upper whisker starts from the hinge and ends at the highest value that is within 1.5 * inter-quartile range of the hinge. The lower whisker extends form the hinge to the lowest value that is within 1.5 * inter-quartile range. The x-axis is consistent between both figures, therefore some homologs (e.g. Glucose–6–phosphate isomerase – directly after UVR irradiation) will have missing data in their specific figure, as there were no differentially expressed *in silico* translated transcripts mapping to that homolog during that time.

In the tricarboxylic acid (TCA) cycle, we observed 14 differentially expressed *in silico* translated transcripts mapping to 5 homologous Uniprot proteins in the TCA cycle (Supplementary Table S12). The rate limiting step of the TCA cycle, isocitrate dehydrogenase, increased in abundance compared to non-irradiated cells, however the variability between individual transcript expression was very large (Fig. 5). Similar to the transcriptomic results observed in the glycolytic pathway, the overall expression of the TCA cycle was mixed with the same number of up and downregulated transcripts (Fig. 5, Supplementary Table S12).

**Figure 5:**
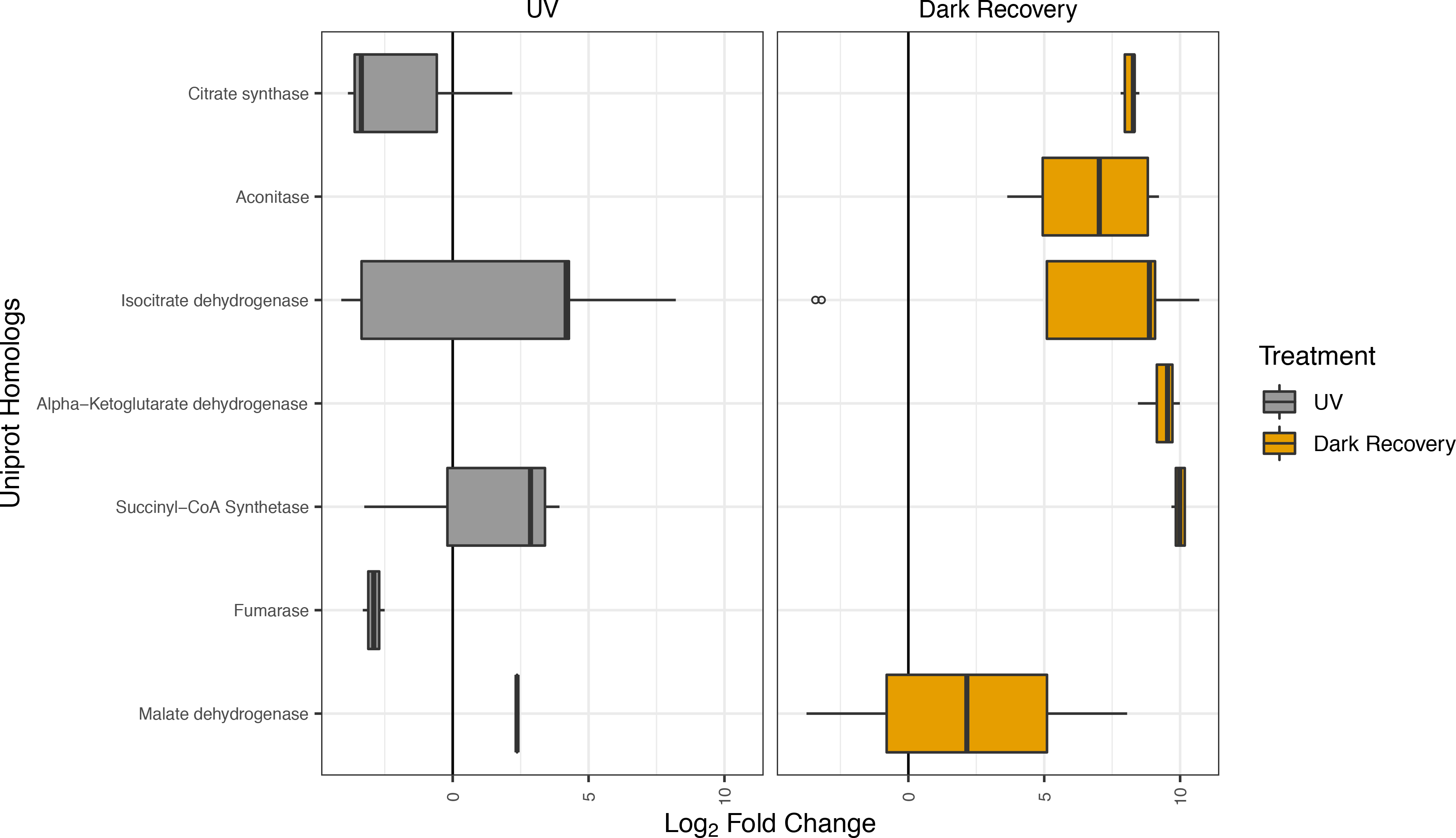
Box and whisker plot for Uniprot TCA homologs. Log_2_ fold changes directly after UVR irradiation are represented in the left figure – labeled “Day” – while log_2_ fold changes after six-hour dark recovery – labeled “Night” – are represented in the figure to the right. The line within the box is the median of the log_2_ fold changes for that specific homolog. The hinges are the 1^st^ and 3^rd^ quartile. The upper whisker starts from the hinge and ends at the highest value that is within 1.5 * inter-quartile range of the hinge. The lower whisker extends form the hinge to the lowest value that is within 1.5 * inter-quartile range. The x-axis is consistent between both figures, therefore some homologs (e.g. Aconitase – directly after UVR irradiation) will have missing data in their specific figure, as there were no differentially expressed *in silico* translated transcripts mapping to that homolog during that time.

### DNA Repair

There were 70 DNA repair transcripts that were differentially expressed after UVR radiation based on ontological mappings (Supplementary Table S13). Out of those transcripts, 64% (45/70) were decreased in abundance compared to the non-irradiated cells (Fig. 6). Transcripts that increased in abundance were related to recombinational DNA repair. Nine *in silico* translated transcripts mapped to *RecA* homologs (Fig. 6), all of which were increased in abundance, including a transcript with a log2 fold change of 8.04 (256-fold increase) – the largest increase in abundance from the DNA repair pathway over non-irradiated cells. There is also an upregulated transcript related to a photolyase homolog, an enzyme that plays a crucial role in UV-induced DNA repair.

**Figure 6:**
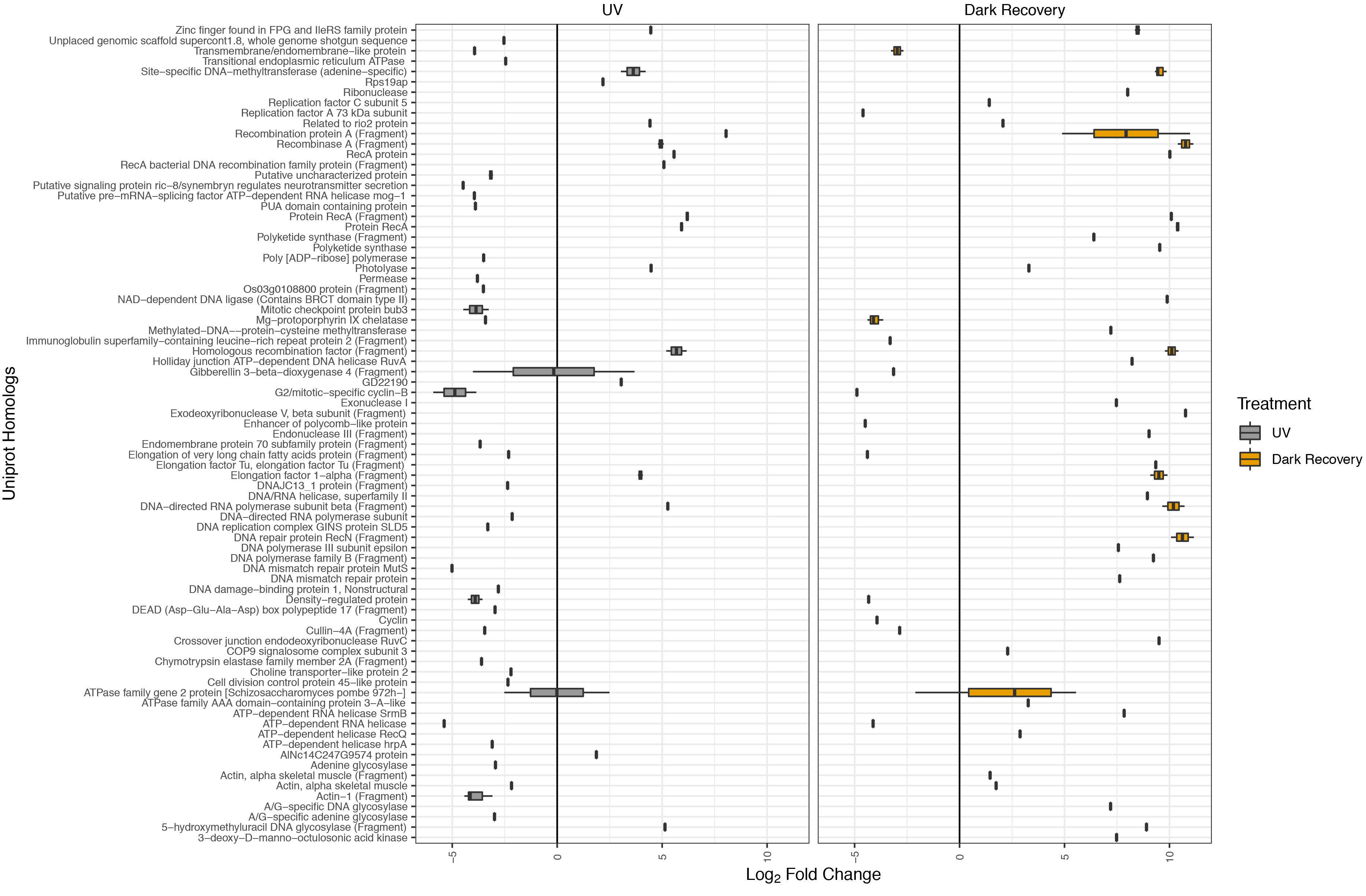
Box and whisker plot for Uniprot DNA repair homologs. Log_2_ fold changes directly after UVR irradiation are represented in the left figure – labeled “Day” – while log_2_ fold changes after six-hour dark recovery – labeled “Night” – are represented in the figure to the right. The line within the box is the median of the log_2_ fold changes for that specific homolog. The hinges are the 1^st^ and 3^rd^ quartile. The upper whisker starts from the hinge and ends at the highest value that is within 1.5 * inter-quartile range of the hinge. The lower whisker extends form the hinge to the lowest value that is within 1.5 * inter-quartile range. The x-axis is consistent between both figures, therefore some homologs will have missing data in their specific figure, as there were no differentially expressed *in silico* translated transcripts mapping to that homolog during that time.

### Preliminary Recovery

Following UVR exposure, we conducted a dark recovery period of 6 hours and measured the transcriptional response. In contrast to the results seen directly after UVR exposure and cellular damage, transcripts in the auxiliary metabolic pathways greatly increased in abundance after dark recovery (Fig. 3-6, Supplementary Tables S14-S17). Approximately 90% of the translated transcripts that mapped to homologous glycolytic and TCA cycle pathway homologs increased their abundance, even after a short dark recovery period of 6 hours. Many transcripts produced log_2_ fold changes greater than two, with several greater than log_2_ of seven. Furthermore, we observed a notable increase in the amount of DNA repair proteins that were upregulated after dark recovery (Fig. 6). After UVR exposure, approximately 36% of the DNA repair transcripts were increased in abundance compared to the non-irradiated cells; however, after dark recovery that number increased to approximately 76% of the transcripts. It was also observed that transcripts that were increased in abundance directly after irradiation seemed to further increase their abundance after dark recovery.

## Discussion

Controlled laboratory studies describing the impact of UVR on phytoplankton have the potential to improve our understanding of photosynthetic damage and repair as well as general DNA damage and repair. To develop a controlled laboratory methodology for studying the impacts of UVR on photosynthetic and metabolic pathways, we developed a UV light emitter to deliver repeatable doses at a high resolution across a wide spectrum of intensities to monitor for break points in photochemical efficiency using FRRF. To test our emitter array, we selected a cosmopolitan open-ocean diatom, of which members of its genus are found in the Southern Ocean where seasonal UV intensity is high due to ozone depletion. For example, during the spring, pennate and centric diatoms comprise a large portion of the Southern Ocean biomass, and the genus *Corethron* is one of the top four genera (Vincent 1988). We chose the diatom *C. hystrix* for our study.

### Growth Rate and Cell Morphology

The growth rate for laboratory grown *C. hystrix* under normal photosynthetically active radiation (PAR, ~40 μmol m^−2^ s^−1^) conditions was similar to published values for several diatoms under analogous light levels (Gilstad and Sakshaug 1990). Higher level plants, which can maintain chlorophyll levels during UVR exposure, may have a higher tolerance to UVR stress as they are able to more efficiently transfer their excitation energy into PSII reaction center proteins (Bornman and Vogelmann 1991; Greenberg et al. 1997). Chloroplast morphology was indeed altered at UVR levels of 0.96-1.59 mW/cm^2^, however the largest decrease in intact chloroplasts occurred mostly during the last two hours of irradiation as result of the cumulative UVR dose. Morphological alterations consisted of jagged chloroplasts that migrated toward the center of the cell, which could be a response similar to the “chloroplast clumping” phenomenon seen in other organisms as a form of UV protection (Sharon et al. 2011).

### Photosynthetic Kinetics and Gene Expression

Photosystem II damage was affected by both UVR intensity and time during our study, with damage from the highest irradiation treatments ultimately resulting in a loss of photosynthetic function – Fv/Fm becomes approximately zero (Fig. 2A, Supplementary Fig. S4-S8). The observed decrease in Fv/Fm is likely caused by UVR directly damaging PSII reaction centers while also increasing *C. hystrix’s* susceptibility to photoinactivation (Rijstenbil 2002). We also observed a corresponding increase in sigma, the functional cross section of PSII which is proportional to the amount of number of chlorophyll molecules per photosystem II reaction center. This increase may be an environmental adaption to the extreme conditions, where under these conditions, chlorophyll molecules may be transferring their excitation energy away from the damaged reaction centers and into the remaining functional reaction centers, thus increasing the amount of chlorophyll molecules servicing the functional centers and increasing the efficiency of energy transfer of those intact centers.

We observed that UVR intensity begins to have a significant effect on the photosynthetic efficiency and energy transfer by 3 hours at the UV intensity of 0.64 mW/cm^2^. These results suggest that the time-dependent damage exceeded repair, provoking an increase in photoinactivation. Furthermore, higher UV intensities (1.28 and 1.59 mW/cm^2^) accelerate photoinactivation much more rapidly, because more reaction centers are being irrevocably damaged. The observed decrease in sigma after 3-4 hours (Fig. 2B), for both 1.28 and 1.59 mW/cm^2^, indicates that the majority of the reaction centers have been damaged or that chlorophyll can no longer funnel the excitation energy through the functional reaction centers (i.e., a loss of photosynthetic function). Thus, it is clear that photosystem operation is maintained at moderate UVR intensities (0.32-0.96 mW/cm^2^) but the cell can only withstand irradiation for several hours before the photosystem is unable to absorb and transfer energy.

For various physiological processes, total dose as compared to dosage rate is important as some changes are the result of a cumulative UVR effect (Frohnmeyer and Staiger 2003; Nawkar et al. 2013). Photosynthetic efficiency (Fv/Fm) decreased twice as fast in cultures irradiated at 1.59 mW/cm^2^ than the culture irradiated at 0.64 mW/cm^2^. Low Fv/Fm measurements represent a decrease in the efficiency of photosynthesis (i.e., dynamic photoinhibition), as seen in the coccolithophorid *Emiliania huxleyi* due to high photon flux densities (Critchley 2000; Ragni et al. 2008). Environmental stressors, such as high intensities of photosynthetically active radiation (PAR), adverse temperatures, and water limitation, typically cause photoinhibition. Our decreased Fv/Fm measurements after UVR exposure appear to mimic photoinhibition, which is possibly due to direct damage to PSII by UVR. However, future applications should determine whether molecular changes under a fast damage rate (high cumulative UV dose and rate) are the same as those changes under a more gradual damage rate (high cumulative UV dose but lower rate).

Photosystem II is known to be the most sensitive part of the photosynthetic system to UVR exposure, especially the oxygen evolving complex (Post et al. 1996; Szilárd et al. 2007). Our transcriptomic analysis shows that oxygen evolving complex genes, *psbO* (PSII manganese-stabilizing protein), *psbP* (PSII oxygen evolving enhancer protein) and psbV (cytochrome c-550), significantly decreased in abundance. *PsbP* has also been discovered to have a supplemental role in stabilizing the PSII-light harvesting super complexes, with decreasing levels of *psbP* producing a concurrent decrease in the amount of super complexes (Ifuku and Noguchi 2016). We observed that the majority of light harvesting complex transcripts decreased in abundance compared to the non-irradiated controls, especially the CP43 and CP47 related transcripts (Ifuku et al. 2011; Ifuku and Noguchi 2016). The CP47 chlorophyll apoprotein along with its sister protein CP43 are functionally used by the cell to channel the energy from the light harvesting complexes into the reaction center core, (Falkowski and Raven 2007). A previous study observed that both CP47 and CP43 protein expression levels decreased in the cyanobacteria *Spirulina platensis* during UVR exposure (Rajagopal et al. 2000). Furthermore, the D1 reaction center protein and the aromatic tyrosine electron donors have been shown to be sensitive to UVR (Vass et al. 1996; Bouchard et al. 2006). D1 is one of two main reaction center proteins, while tyrosine amino acids are part of the donor side of PSII absorbing energy at 285 nm, thus making them a possible target for UVR irradiation (Vass et al. 1996). Transcriptome changes demonstrate that both the D1 and D2 reaction center proteins, encoded by *psbA* and *psbB* respectively, mapped to transcripts which decreased in abundance during UVR exposure compared to non-irradiated cells. It has been determined that the overall changes in the expression of *psbA* differ based on the specific organism (Surplus et al. 1998; Huang et al. 2002), however, the decrease in abundance of our *psbA* transcripts during UVR was similar to the changes produced by *Arabidopsis thaliana* (Surplus et al. 1998). Other PSII reaction center transcripts, such as those that mapped to the cytochrome b559 protein, also displayed large decreases in abundance after UVR exposure.

Intuitively, it would be assumed that if these subunits were being actively damaged there would be an increase in their individual gene expression to help mitigate and replace damaged protein subunits. UVR exposure, however, has many other cellular consequences including an increased amount oxidative stress within the cell (Rijstenbil 2002). The Reactive Oxygen Species (ROS) produced during oxidative stress can have serious deleterious effects including photodamage to photosynthetic machinery, an imbalance in the photosynthetic redox signaling pathways and direct inhibition of D1 synthesis, which is necessary for PSII repair (Gururani et al. 2015). The result of these effects is increased photosynthetic photoinhibition. After UVR irradiation, we observed an upregulation of transcripts related to ATP-dependent Clp proteases and cysteine proteases, chloroplast specific enzymes which play a role in protein turnover, suggesting the cells may indeed be dealing with an increased amount of damaged proteins (Olinares et al. 2011). Additionally, ROS can also activate a signaling cascade, where receptors transmit signals to regulatory molecules which decrease nuclear gene expression of photosynthetic genes (Gururani et al. 2015). This likely explains why nearly complete downregulation of photosynthetic genes is observed for organisms subjected to UVR (Rijstenbil 2002; Gururani et al. 2015).

Tau is a proxy for the PSII reaction center turnover time (Kolber et al. 1988). Both tau 1 and tau 2 turnover times increase significantly at UVR intensities of 0.64-1.59 mW/cm^2^ (Fig. 2CD). This is because the plastoquinones QA and QB are directly susceptible to UVR damage (Melis et al. 1992). Our data also illustrate that turnover time corresponds to intensity (Fig. 2C-D, Supplementary Fig. S4-S8, Supplementary Tables S3-S7, S9 and S10). With turnover times increasing up to 0.14 seconds for tau 2 at 1.28 mW/cm^2^ (Fig. 2D), these long reoxidation times have a detrimental effect on photosynthesis as excess photon energy causes a buildup of oxidative radicals, thus explaining the genetic downregulation of photosynthetic genes and almost complete loss of photosynthetic function at higher intensities of UVR.

Previously it was observed that the redox state of QA may have an effect on the abundance of light harvesting complex transcript abundances (Maxwell et al. 1995). This phenomenon is perhaps another driving force behind the decrease in abundance in the majority of light harvesting complex transcripts. As the turnover time for QA increases during UVR exposure, the cell may reduce the abundance of certain light harvesting proteins in order to prevent an increase in irradiance stress and higher photoinhibition. Simultaneously, the cell maybe trying to maintain the structural integrity of functional reaction centers (through an increase in chlorophyll binding protein), as a way of limiting damage. This type of competing self-regulation highlights the overall complexity of transcriptomic regulation during UVR exposure.

Transcriptome analysis of supplemental photosynthetic electron transport genes corresponds with our PSII expression analysis, as the majority of the transcripts were downregulated compared to the control samples. Most of the transcripts in this subset of data mapped to homologous cytochrome b6-f complex proteins. There are 4 major subunits that make up the cytochrome b6-f complex in algae (Pierre et al. 1995). We identified both of the heme-bearing subunits of the cytochrome b6-f complex, and they were strongly downregulated after UVR exposure. Thus, the cell extends its protective strategy to UVR exposure, using transcriptional downregulation, to complex III in the photosynthetic electron transport chain.

Photosystem I is not directly affected by UVR and is more resistant to environmental stressors such as high-light levels (Teramura and Ziska 1996; Zhang et al. 2016). However, with an interaction between PSII and PSI due to the photosynthetic electron cascade, UVR exposure will unavoidably have a downstream impact on photosystem I. The transcriptional analysis provided some insight into the health of photosystem I. Of the 16 transcripts from photosystem I, 15 transcripts mapping to four Uniprot homologs, decreased in abundance during irradiation. The majority of the PSI transcripts map to *psaA* and *psaB* which code for PSI P700 chlorophyll *a* apoprotein A1 and A2 respectively. *PsaB* binds hydrophobically to *psaA* to form the major reaction center of PSI (Falkowski and Raven 2007). Additionally, the PSI subunit II (*psaD*) transcripts were also significantly downregulated. *PsaD* is an important protein that is crucial for the stability and correct assembly of PSI while also providing a ferredoxin binding site (Hou et al. 2017). The decreased abundance of *psaA*, *psaB*, and *psaD* transcripts indicates that like PSII and the b_6_f complex, transcriptional downregulation was most likely influenced by PSII photoinhibition and oxidative stress, which means it could possibly act as a secondary protective strategy to prevent increased stress and photoinhibition.

The first stage of the photosynthetic dark reactions is the fixation of CO_2_ into 3-phosphoglycerate, which is catalyzed by the protein RuBisCO. RuBisCO is the most abundant protein on Earth due to its crucial function combined with its inefficiency as a carboxylase (Raven 2009; Raven 2013). Our data demonstrated a coordinated downregulation for the transcripts mapping to the RuBisCO homologs, with all transcripts decreased in abundance compared to non-irradiated cells (Fig. 3). Previous research on higher order plants demonstrated that UVR exposure caused a large decrease in the activity and expression of RuBisCO when compared to control samples in pea leaves (Strid et al. 1990; Mackerness et al. 1999). Moreover, similar RuBisCO decreases were also observed in jackbean leaves (Choi and Roh 2003). This implies that the mechanism for gene expression decreases in RuBisCO expression ties directly back to the activity of the RuBisCO protein as well as through direct deactivation of its synthesis, similar to these previous studies (Choi and Roh 2003).

### Metabolic Pathway Expression

UVR exposure is known to cause other metabolic gene expression changes in higher level plants (Jenkins 2009). The glycolytic cycle is the first phase in the catabolism of cellular carbohydrates (Nelson and Cox 2005). The TCA cycle, the second phase of catabolism, is an important source of the cellular reducing agent NADH, which helps generate the proton gradient that is critical for the production of ATP through electron transport (Nelson and Cox 2005). In our study, a coordinated downregulation of transcripts mapping to glycolysis and the TCA cycle was not observed, rather we observed a mixed gene expression pattern directly after UVR exposure (Fig. 4, 5), suggesting that certain enzymes may be shared between multiple pathways for other purposes. Logemann et al. 2000 found that UV-induction of select primary metabolism enzymes that can provide carbon substrates for the shikimate pathway, while Casati et al. 2003 discovered that certain primary metabolism enzymes may be induced to provide energy in the form of ATP for the synthesis of molecules necessary for cell survival under UVB stress (Logemann et al. 2000; Casati and Walbot 2003).

### DNA Repair

Diatoms are subjected to high doses of UVR in their natural environment (Karentz et al. 1991; Fricke et al. 2011), and the selective pressure of managing UV damage may provide novel insights into mechanisms of DNA repair. Because DNA can directly absorb UVR, the damaged DNA forms cyclobutane pyrimidine dimers (CPDs) or pyrimidine 6-4 photoproducts (6-4PPs) (Sinha and Häder 2002). These photoproducts (i.e., lesions) can cause DNA to make an unnatural conformation, arresting replication (Rastogi et al. 2010). To minimize mutagenic effects, several organisms, including diatoms, produce photoreactivating proteins called photolyases which directly remove the deleterious dimers (Coesel et al. 2009). Our data demonstrate that a single transcript mapped to a photolyase homolog, and it significantly increased in abundance directly after irradiation (Fig. 6). With its important role in DNA repair, future studies should identify the functional significance for this particular photolyase and how it affects the overall fitness of *C. hystrix* during UVR exposure.

Additionally, there are times where these photoproducts can lead to secondary DNA breaks, or UVR is so intense that double stranded breaks become one of the main photochemical end-products (De Mora et al. 2000). With the emergence of double-stranded breaks, the gene *recA* is recruited to initiate the cellular SOS response, which regulates between 50 and 66 genes involved in double stranded DNA (dsDNA) repair (Smith et al. 1987; Janion et al. 2002). *RecA* mediates homologous recombination repair, which will function to maintain the integrity of DNA. We observed all transcripts mapping to Uniprot *recA* homologs significantly increased in abundance relative to non-irradiated cells (Fig. 6), implicating the activity of these enzymes in dsDNA repair during our study. Furthermore, two site specific DNA-methyltransferase transcripts, which are known to induce the cellular SOS response, were also significantly increased in abundance directly after UVR exposure (Heitman and Model 1987) further highlighting the cells adaption to managing UVR exposure.

### Preliminary Recovery

After 6 hours of recovery in dark conditions, several photosynthetic light-harvesting transcripts increased in abundance compared to the non-irradiated samples, with some homologs demonstrating a reversal in expression (Fig. 3, Supplementary Table S14). These results suggest that the cell is possibly trying to repair certain aspects of its UV damaged photosynthetic clusters in order to survive, as photodestruction can cause a complete loss of function for the complex (Lao and Glazer 1996). As the light harvesting complex provides the light energy for the photosystem reaction centers, it would be beneficial for the cell to repair the harvesting complex. Furthermore, homologous transcripts that were part of the cytochrome b6/f complex or maintained roles in photosynthetic electron transport also increased in abundance after night recovery. The largest change in abundance was by a *petF* transcript, which codes for the photosynthetic ferredoxin protein. This expression reversal may indicate that the cells are trying to restart and repair electron transport before PAR light returns to re-energize the photosynthetic system, or that ferredoxin may have a supplemental role in the defense against lingering oxidative stress similar to its response after other biotic attacks (Bilgin et al. 2010).

Moreover, despite the tremendous diversity of diatoms, with species estimates ranging from 1 × 10^4^ (Norton et al. 1996) to 2 × 10^5^ (Allen et al. 2006), few studies have focused on the potential biomedical applications of bioactive compounds produced by these organisms (Coesel et al. 2009; Prestegard et al. 2009). There were two polyketide synthetase transcripts which were significantly upregulated after dark recovery (Fig. 6). These enzymes are important in the biosynthesis of many natural products and could be involved in the production of compounds such as mycosporine-like amino acids (MAAs) (Klisch 2008).

Still, while there was an upregulation of some transcripts during the dark recovery period, many transcripts especially PSI and II transcripts, were still downregulated after dark recovery (Supplementary Table S14). These results suggest that while the cell is actively trying to repair its photosystem, the damage may have been severe enough that recovery of the whole photosynthetic pathway was not possible during the six-hour recovery period (Neale et al. 1998; Fritz et al. 2008).

Our emitter array produced morphological, physiological and molecular responses with extremely high resolution and reproducibility. We demonstrate, based on *C. hystrix* physiological measurements, that our emitter array allowed us to directly manipulate the rate and intensity of photosynthetic damage based on the specific applied UVR energy intensities. We were also able to modulate metabolic gene expression changes. This ability to study specific physiological and molecular responses over a large spectrum of UVR intensities could provide further insights into UV induced damage in other complex organisms. Large-scale screening of organisms that are well adapted to high UV fluxes may contain novel mechanisms or natural compounds, such as photolyases or MAAs, which could increase our understanding of preventive mechanisms and possible treatments resulting from DNA damage.

## Acknowledgements

We would like to thank Dr. Zbigniew Kolber for his expertise and guidance during the manuscript preparation.

## Compliance with Ethical Standards

### Conflict of interest

Authors Robert Read, David Vuono and Iva Neveux declare that they have no conflict of interest. Authors Joseph Grzymski and Carl Staub have a conflict of interest as they are co-founders of the company, EMS Genomics, that developed the light engine.

### Ethical approval

This article does not contain any studies with human participants or animals performed by any of the authors.

